# Single-nucleotide resolution profiling reveals dynamic and site-specific m⁶A regulation during human myogenesis

**DOI:** 10.64898/2026.01.09.697937

**Authors:** Pierre Klein, Krizia Ronquillo, Ludovic Arandel, Frédérique Rau, Denis Furling

## Abstract

**Background:** Myogenic differentiation of muscle stem cells required for skeletal muscle growth and regeneration relies on the precise spatiotemporal coordination of multiple gene-expression programs. Over the past decades, considerable progress has been made in defining the transcriptional networks that govern myogenesis. In particular, myogenic transcription factors integrate extrinsic and intrinsic cues to drive skeletal muscle development and regeneration. In contrast, post-transcriptional layers of regulation have remained comparatively underexplored. Among these, the epitranscriptome – comprising more than 170 chemical RNA modifications – has recently emerged as a major regulatory axis in cellular differentiation. N⁶-methyladenosine (m⁶A), the most abundant internal modification of mRNA, is now recognized as a key regulator of stem-cell fate decisions across multiple biological systems. By dynamically modulating RNA stability, translation, and other processing steps, m⁶A enables precise fine-tuning of gene-expression programs in response to physiological cues. These effects are mediated through m⁶A-binding RNA-binding proteins (RBPs), also referred to as “readers”, which selectively recognize methylated transcripts and translate the modification into functional cellular outcomes. Despite its central role in other tissue contexts, the contribution of m⁶A to skeletal muscle physiology – and particularly to human muscle stem cell biology – remains poorly characterized. This gap represents a critical and largely unexplored frontier in muscle biology.

**Methods:** To define the epitranscriptome landscape during the myogenic differentiation of human skeletal muscle cells, we performed GLORI on proliferating and differentiated human primary myoblasts. This recently developed approach enables detection of m⁶A at single-nucleotide resolution together with quantitative measurement of methylation stoichiometry at individual sites. m⁶A-modified positions were mapped across the transcriptome in both cellular states and integrated with matched RNA-seq datasets data to contextualise m⁶A features relative to transcript abundance. To place these findings in a broader context, we compared our human m⁶A maps with previously published mouse muscle cell datasets and overlaid them with available CLIP-seq datasets of m⁶A RNA-binding proteins.

**Results:** We identified tens of thousands of high-confidence m⁶A sites in proliferating and differentiated human myogenic cells. While many sites were shared between states, myogenic differentiation was accompanied by epitranscriptomic remodeling, affecting both the distribution and stoichiometry of m⁶A marks. Transcripts exhibiting differential m⁶A regulation were enriched for pathways related to myogenic development, cytoskeletal remodeling, signalling, nucleic-acid–associated processes and energy metabolism. Joint analysis of m⁶A site number, methylation stoichiometry, and expression levels indicated the presence of distinct m⁶A regulatory frameworks at the transcript level. Genes associated with myogenesis and regulatory pathways were distributed across these categories, consistent with the coexistence of multiple regulatory programs during myogenic differentiation. In addition, cross-species comparisons revealed the presence of human-specific methylation patterns. Integration with published m⁶A RNA-binding protein datasets uncovered distinct subsets of m⁶A sites with varying binding profiles, suggesting state-dependent interpretation of m⁶A marks.

Together, these data constitute, to our knowledge, the first single-nucleotide–resolution atlas of m⁶A methylation in human skeletal muscle, revealing a previously uncharacterized epitranscriptomic landscape and providing a foundational resource for studies of human myogenesis.

## Background

Skeletal muscle formation and regeneration is a complex and highly coordinated process that requires tight spatiotemporal control of gene-expression programs governing progenitor maintenance, lineage commitment, and terminal differentiation [1]. Decades of research have defined the transcriptional networks that orchestrate myogenesis, establishing myogenic transcription factors as central integrators of intrinsic and extrinsic cues [2]. By contrast, post-transcriptional regulatory layers — which determine how, when, and where transcripts are processed and translated — remain disproportionately understudied in skeletal muscle biology [3,4].

Within this broader regulatory landscape, RNA modifications – collectively referred to as the epitranscriptome and comprising more than 170 chemical modifications – have emerged as a dynamic and versatile layer that modulates RNA fate and function [5,6]. The most abundant internal modification in eukaryotic mRNA, N⁶-methyladenosine (m⁶A), is deposited by the METTL3–METTL14–WTAP methyltransferase complex at RRACH motifs and influences multiple aspects of the mRNA life cycle, including splicing, stability, translation, and decay. Through these actions, m⁶A plays key roles in cell differentiation, lineage commitment, developmental transitions, and metabolic homeostasis. Profiling studies across diverse systems have established m⁶A as a pervasive regulator of gene-expression programs, with thousands of human transcripts carrying this modification [7].

A defining feature of m⁶A-mediated regulation is its ability to control stem-cell fate through subtle, site-specific modulation of select transcripts rather than wholesale changes in global methylation. In embryonic stem cells, for example, a limited subset of transcripts undergoes cell-type–specific m⁶A remodeling, yet these changes are sufficient to drive lineage commitment [8–11]. Similar principles have been observed in hematopoietic and neural stem-cell systems [12–15], underscoring m⁶A as a context-dependent post-transcriptional regulator of differentiation.

In skeletal muscle, emerging evidence indicates that m⁶A contributes to myogenic progression, tissue homeostasis, and the response to physiological stress or injury in mouse models. Thus, global m⁶A levels increase following acute muscle damage, concomitantly with the activation and proliferation of muscle stem cell (MuSC) during the regenerative process [16] and decline when myoblasts undergo differentiation as observed *in vitro* using the C2C12 mouse myoblast cell line [16]. Genetic perturbation of m⁶A writers, erasers, or readers affects MuSC proliferation, differentiation, and regenerative capacity [17–20]. However, these findings – largely derived from bulk or antibody-based approaches – have also revealed apparent contradictions, suggesting that the functional outcome of m⁶A signaling depends on precise, site-specific regulation and engagement of m⁶A-binding proteins, rather than on global methylation levels alone.

Despite these insights, the global m⁶A landscape in human skeletal muscle – and its remodeling during human myogenesis – remains unexplored. Existing datasets are limited to non-human models. Moreover, they predominantly rely on antibody-based approaches such as MeRIP-seq, which lack single-nucleotide resolution and do not quantify methylation stoichiometry, and are associated with substantial technical biases and variability [17,19,21]. As a result, the fine-scale regulatory logic by which m⁶A shapes RNA fate and gene-expression programs in human muscle cells remains unknown.

To address this gap, we generated the first single-nucleotide–resolution atlas of m⁶A methylation during human myogenic differentiation using GLORI, a non-biased approach that enables stoichiometry-resolved mapping of m⁶A at individual nucleotides [6]. Applying this strategy to primary human muscle cells undergoing myogenic differentiation, we integrated these high-resolution m⁶A maps with transcript abundance, functional enrichment analyses, datasets from mouse m⁶A studies, and binding profiles of m⁶A readers. This integrated framework provides a comprehensive view of epitranscriptomic regulation in human skeletal muscle and reveals coordinated, state-specific m⁶A programs linked to myogenic fate transitions.

## Method

### Tissue Culture

Primary human myoblasts (CHQ, gift from V. Mouly) were derived from the quadriceps muscle of a 5-day-old female biopsies, as previously described [22,23]

Proliferating myoblasts were maintained in a growth medium consisting of 64% DMEM, GlutaMAX™ Supplement (Gibco, cat. no.31966) and 16% 199, GlutaMAX™ Supplement (Gibco, cat. no.41150) media supplemented with 20% fetal bovine serum (FBS, SERANA cat. no.S-FBS-CO-015), 5 µg/mL insulin (Sigma Aldrich, cat. no.910776), 25 µg/mL fetuin, 5 ng/mL human epidermal growth factor (Gibco, cat. no.PHG0311), and 0.5 ng/mL basic fibroblast growth factor (Gibco, cat. no.PHG0026). Cultures were kept at 37 °C in a humidified 5% CO₂ incubator and passaged at low confluence to preserve proliferative capacity.

Myogenic differentiation was initiated when cultures reached high confluence (∼95%), by replacing the growth medium with serum-free DMEM, GlutaMAX™ Supplement (Gibco, cat. no.31966). Medium was renewed every 48 h, and cells were collected after 7 days of differentiation.

HEK293T cells were maintained at 37 C in DMEM, GlutaMAX™ Supplement (Gibco, cat. no.31966) supplemented with 10% fetal bovine serum (FBS, SERANA cat. no.S-FBS-CO-015), and 1% penicillin–streptomycin (Gibco, cat. no.15070063) in a humidified 5% CO₂ incubator.

### Total RNA extraction

For each condition (proliferating or differentiated myoblasts and HEK293T cells), biological replicates (cells grown on two 150-cm² plates) were pooled prior to RNA extraction to obtain sufficient material. Cells were washed twice with PBS 1× to remove serum traces, detached using trypsin–EDTA (Gibco), and pelleted by centrifugation (300 × g, 5 min at RT). Pellets were washed once in ice-cold PBS, centrifuged again, and snap-frozen on dry ice.

Total RNA was extracted using TRI Reagent (Sigma-Aldrich, T9424) according to the manufacturer’s protocol, including the chloroform phase-separation step, isopropanol precipitation, and ethanol washes. RNA pellets were resuspended in nuclease-free water. RNA quantity and purity (A260/A280 and A260/A230 ratios) were assessed using a NanoDrop spectrophotometer (Thermo Scientific™ NanoDrop™ One).

### Poly(A) enrichment

Polyadenylated RNA was isolated using Dynabeads Oligo(dT)25 (Thermo Fisher, 61005) following the manufacturer’s instructions. Briefly, for each sample, 75 µg of high-quality total RNA was used as input. RNA was denatured at 65 °C for 2 min in an equal volume of binding buffer and immediately placed on ice to disrupt secondary structures. The denatured RNA was then incubated for 5 min with pre-equilibrated oligo(dT)25 beads under gentle head-to-tail rotation to allow hybridization of poly(A) tails. Beads were washed multiple times with the high-salt wash buffers (10 mM Tris-HCl, pH 7.5, 0.15 M LiCl, 1 mM EDTA, 0.1% LiDS) to remove rRNA and other non-polyadenylated transcripts. Poly(A)⁺ RNA was eluted by heating the bead–RNA complexes at 80 °C for 2 min in nuclease-free water. The eluate was subsequently subjected to a second independent purification round, using freshly regenerated oligo(dT)25 beads, to further increase the purity of the polyadenylated fraction. The final Poly(A)⁺ RNA eluate in RNAse free water, quantified using NanoDrop spectrophotometer (Thermo Scientific™ NanoDrop™ One) and stored at −80 °C until further processing.

### GLORI Treatment

Poly(A)⁺ RNA was subjected to the GLORI method (glyoxal- and nitrite-mediated deamination of unmethylated adenosines to inosines) to detect m⁶A residues at single-nucleotide resolution, with each CHQ sample processed in technical replicate, as described in the original protocol [6]. Briefly, 200 ng of poly(A)⁺ RNA per reaction was fragmented at 94 °C for 4 min in fragmentation buffer (New England BioLabs, cat. no. E6186), immediately cooled on ice to prevent over-fragmentation, and supplemented with 2 µL Stop Solution (New England BioLabs, cat. no. E6187). The fragmented RNA was then ethanol-precipitated and resuspended for subsequent GLORI processing. Guanosines in the fragmented RNA were protected using a buffer containing 1.32 M glyoxal solution (Sigma-Aldrich, 50649) and 50% DMSO (Sigma-Aldrich, red D8418) in nuclease-free water. The mixture was incubated in a preheated thermocycler at 50 °C for 30 min. After incubation, tubes were returned to room temperature and supplemented with 10 µL saturated H₃BO₃ (Sigma-Aldrich, B0394), followed by an additional 30 min incubation at 50 °C. Next, a deamination buffer consisting of 1.5 M NaNO₂ (Sigma-Aldrich, 31443), 80 mM MES (pH 6.0), and 1.76 M glyoxal in RNase-free water was added, and samples were incubated at 16 °C for 8 h to selectively deaminate unmethylated adenosines. RNA was purified by ethanol precipitation. Finally, guanosines were deprotected by adding a deprotection buffer (500 mM TEAA and 45% deionized formamide) and incubating at 95 °C for 10 min. RNA was again purified by ethanol precipitation.

### Library Preparation

GLORI-treated RNA was then used for library preparation. Deprotected RNA was first repurified using the Clean & Concentrator-5 kit (Zymo Research, red R1013) according to the manufacturer’s instructions. Purified RNA was dephosphorylated with 0.5 units of T4 Polynucleotide Kinase (NEB, M0201S) at 37°C for 1h, then at 65 °C for 20 min. Following ethanol precipitation, 1 µL of 20μM 3′ linker (5′rAPP-AGATCGGAAGAGCGTCGTG-3SpC3) was ligated using 10 units of T4 RNA Ligase 2, truncated KQ (NEB, M0373S) at 25 °C for 2h. Excess adapter was removed enzymatically using 50 units of 5′ deadenylase (NEB, M0331S) and 30 units of RecJf (NEB, M024S), upon incubations at 30 °C and 37 °C for 1 h respectively, followed by heat inactivation at 70°C for 20 min.

RNA was purified by ethanol precipitation and reverse-transcribed in a buffer containing 1 µL of 2 µM reverse transcription primer (ACACGACGCTCTTCCGATCT) using 10 units of SuperScript III (Invitrogen, 18080093). Excess primer was removed by incubation with 20 units of exonuclease I (NEB, M0293S) at 37°C for 30 min.The fractions was stopped by adding 1ul of 0.5M EDTA (pH 8.0) (Thermo Fisher Scientific, cat. no. R1021). RNA–cDNA hybrids were degraded under strong alkaline conditions, and the cDNA was purified using MyOne Silane beads (Thermo Fisher Scientific, cat. no. 37002D).

Next, 0.8 µL of 80µM 5′ adapter (5′Phos-NNNNNNNNNNAGATCGGAAGAGCACACGTCTG-3SpC3) containing a 10-nt UMI was ligated directly on beads using 0.77 units T4 RNA Ligase 1, high-concentration (NEB, M0437M) with 1.02 mM of ATP (New England BioLabs, cat. no. N0437). cDNA was purified again using MyOne Silane beads and amplified by PCR using 1x NEBNext Ultra II Q5 Master Mix (New England BioLabs, cat. no. M0544S), 0.2 µM NEBNext Universal Index Primer of Illumina (New England BioLabs, cat. no. E6609S) and NEBNext Universal PCR Primer for Illumina for 12 cycles.

PCR products were purified using 1.8× AMPure XP beads (Beckman Coulter, cat. no. A63880). Fragments 165–250 bp were size-selected by 8% native polyacrylamide gel electrophoresis. Gel slices were fragmented using pipette tips for 5 min in 400 µL 1× TE buffer, incubated at 37 °C for 1 h at 1,200 rpm, snap-frozen in liquid nitrogen, and incubated overnight at 37 °C with shaking (1,200 rpm). Samples were centrifuged at 14,000×g for 5 min at 4 °C to elute DNA through Spin-X centrifuge tube filter (Costar, cat. no. 8162) The filtrate was ethanol-precipitated.

Final DNA libraries were resuspended and quantified using the Qubit dsDNA High Sensitivity Assay Kit (Invitrogen, cat. no. Q33230).

A control library from untreated RNA was prepared in parallel from 200 ng of poly(A) RNA, following the same procedure as for GLORI-treated samples, except that no deamination buffer was added.

### Sequencing

Indexed libraries were pooled equimolarly and sequenced on an Illumina platform (NovaSeq X) using, paired-end 150-bp reads (PE150) and aiming for 40–50 Gb of raw data per sample, ensuring sufficient depth for quantitative m⁶A stoichiometry analysis. Sequencing was performed by the Biomics Sequencing Core Facility at Institut Pasteur, Paris, France with standard Illumina quality control metrics applied.

### Computational Analysis of GLORI data

#### Calling m6A sites

Sequencing data generated from GLORI-treated libraries were converted from BCL to FASTQ format using bcl-convert (v2.20.0) with default parameters. Adapter and low-quality bases were trimmed using Trim Galore! (v0.6.5; which wraps Cutadapt) with a Phred quality cutoff of 20 and stringency 1, discarding reads shorter than 35 nucleotides. PCR duplicates were removed in a sequence-based manner using seqkit rmdup (v2.9.0, -s option). The 5′ 10-nt UMI was then removed from each read using fastx_trimmer (v0.0.14; -Q 33 -f 11). Control (untreated) libraries were processed using the same preprocessing pipeline. Cleaned reads were processed to identify m⁶A sites using GLORI-tools v1.0, a dedicated bioinformatics pipeline for absolute m⁶A quantification at single-nucleotide resolution. The pipeline is openly available on GitHub from the Luicongcas laboratory. Reference files for GLORI analysis were prepared as follow, annotations for the human reference genome (GRCh38) were downloaded from NCBI, and chromosome names in the GTF file were unified with the genome FASTA using the assembly_report.txt correction script provided with GLORI-tools. Modified reference genomes were generated using build_genome_index.py, which produces the A-to-G–converted forward and reverse complementary genomes required for GLORI alignment. STAR (v2.7.11) indexes were then built separately for each A-to-G–converted genome. A transcriptome reference was generated using gtf2anno.py and build_transcriptome_index.py, producing an A-to-G–converted transcriptome based on the longest transcript per gene from RefSeq/NCBI. The corresponding transcriptome index was built using bowtie v1.3.1.

Single-base annotation tables were created using anno_to_base.py, gtf2genelist.py, and anno_to_base_remove_redundance_v1.0.py, yielding non-redundant base-level annotations for downstream quantification. Cleaned reads were aligned and m6A sites were called and quantified using run_GLORI.py, based on transcriptome-wide per-site A-to-G conversion rates. Reads from untreated control libraries were aligned using the same GLORI alignment workflow provided.

For genome browser visualization and downstream analyses, per-site GLORI methylation tables were converted into standard six-column BED files using a custom R script. Genomic coordinates, gene names, and strand information were retained, and GLORI methylation values were scaled to a 0–1000 BED score. BED entries were sorted by genomic position before use in overlap analyses.

#### Metagene

Metagene profiles were generated from m⁶A sites identified in the totalm6A.FDR.csv files, which were converted to BED format using a custom R script and processed using the MetaPlotR pipeline as previously described [24].

#### Visualization of m⁶A sites

GLORI-aligned BAM files were loaded into the Integrative Genomics Viewer (IGV) alongside reference gene annotations to inspect the distribution of modified positions along transcripts.

#### Motif discovery

Flanking sequences (3 nucleotides in each direction) of each GLORI-identified m6A site from the reference genome were extracted for motif analysis. The motifs logos obtained via MEME (v.4.11) [25] analysis were plotted.

#### Gene Ontology

GO analysis was performed using PANTHER (Protein ANalysis THrough Evolutionary Relationships) v16.0 [26] (http://geneontology.org) and the PAN-GO functionome v2.0 [27], restricting the analysis to the Biological Process category and Homo sapiens. When required, redundant GO terms were removed using REVIGO [28], allowing small or medium term similarity. The top 10 GO terms ranked by fold enrichment were retained, and results were hierarchically ordered by FDR.

#### Reanalysis of publicly available iCLIP data of m6A readers

iCLIP datasets for endogenous YTH family proteins (YTHDF1, YTHDF2, YTHDF3, YTHDC1, YTHDC2) in HEK293T cells [29] were downloaded and reprocessed. Raw bedGraph signal files were lifted from hg19 to hg38 using UCSC LiftOver and processed to retain intervals with uTPM ≥ 1. GLORI-derived single-nucleotide m⁶A sites were expanded to a ±10 bp window and intersected with each reader’s iCLIP signal. A site was considered bound when overlapping enriched iCLIP signal was detected, and all overlapping readers were recorded (multi-binding allowed). For each GLORI site class (proliferation-only, differentiation-only, shared), we quantified the number and percentage of sites bound by each reader and the fraction of sites unbound by any reader. These frequencies were computed relative to the total number of GLORI sites per class.

#### Reanalysis of publicly available MeRIP-seq data in mouse muscle cells

m⁶A peak calls from Tan *et al*. (Supplementary Data 4) [30] for proliferating and differentiated GFP-transfected C2C12 cells were extracted from the provided Excel file, converted to BED format, and lifted from the mouse genome (mm10/GRCm38) to the human genome (hg38/GRCh38) using UCSC LiftOver. These lifted peaks were then overlapped with our GLORI single-nucleotide m⁶A sites to identify conserved m⁶A-modified regions. Mouse gene names were cleaned and converted to their human orthologues using Ensembl Compara via the biomaRt getLDS() function, restricting to one-to-one symbol-level mappings. Only one-to-one orthologues were retained. The resulting human gene lists were then intersected with our GLORI-defined m⁶A-modified genes in proliferating and differentiated human myoblasts to assess conservation of m⁶A regulation between mouse and human myogenesis

### Data analysis

The bioinformatics analyses were performed on the HPC facilities of the National Network of Computing Resources of the Institut Français de Bioinformatique (IFB), funded by the Programme d’Investissements d’Avenir (PIA), grant Agence Nationale de la Recherche number ANR-11-INBS-0013. Data analyses were also conducted using Microsoft Excel (v16.103), GraphPad Prism (v10), and RStudio (v4.5), which were additionally used for data visualization and graph generation. Illustrations were created with BioRender.com, under a Creative Commons Attribution-NonCommercial-NoDerivatives 4.0 International license.

## Results

### Transcriptome-wide identification of m⁶A during human myogenic differentiation

To identify m⁶A marks and their dynamic during myogenic differentiation of human skeletal muscle cells, we performed GLORI on polyadenylated RNA isolated from primary human myoblasts in proliferating and differentiated stages. GLORI enables glyoxal- and nitrite-mediated deamination of unmethylated adenosines, allowing direct single-nucleotide identification of m⁶A residues and quantitative estimation of stoichiometric changes at each site (Fig. 1A). Cellular morphology and expression profiles were consistent with myogenic cultures in both conditions, with high expression of the myogenic marker desmin and quasi-nul expression of the fibroblast marker *PDGFRA* (Fig. 1B).

**Figure 1.**
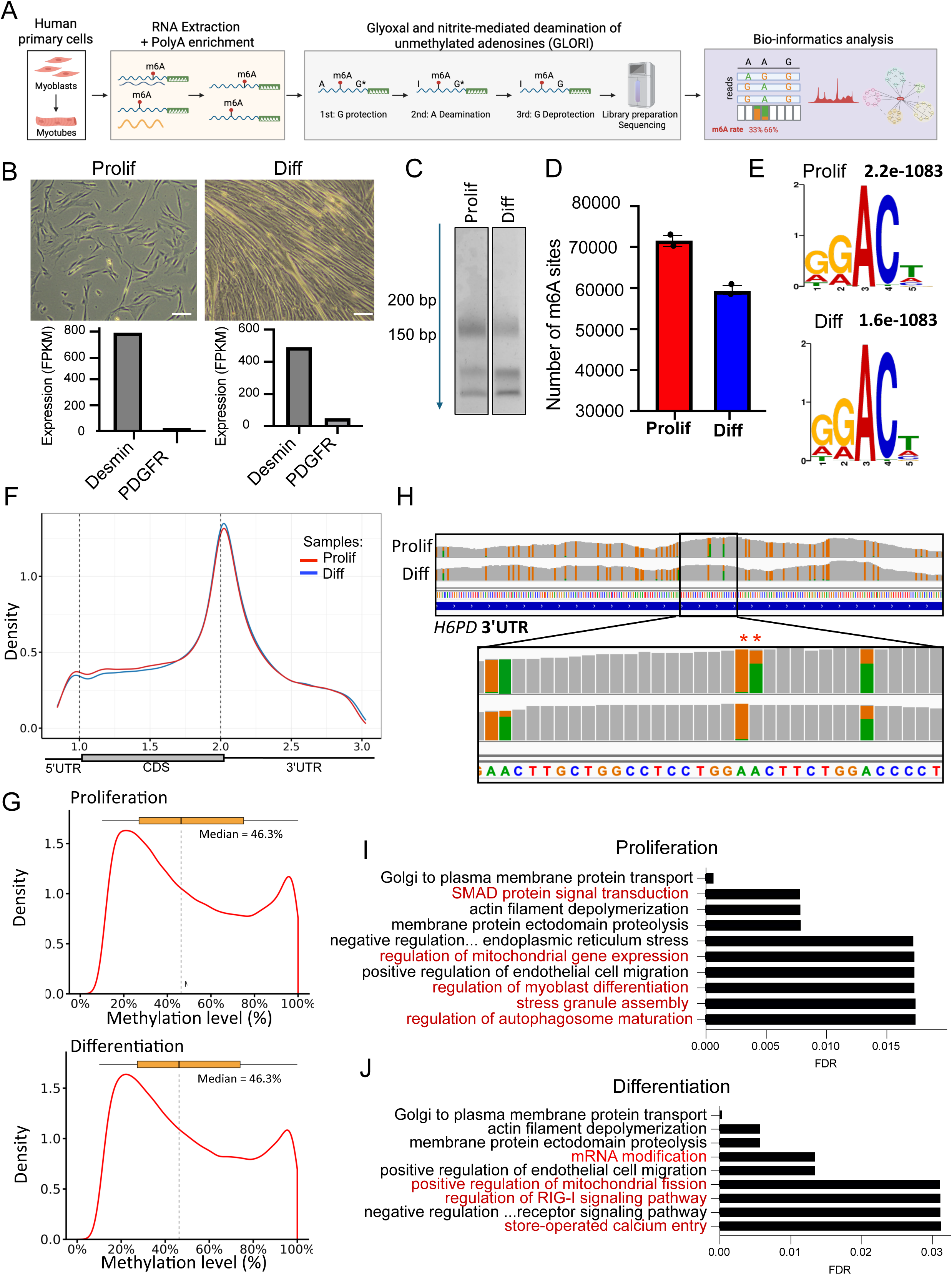
Global landscape and dynamics of m⁶A methylation during human myogenic differentiation. **(A)** Overview of the GLORI workflow applied to primary human myoblasts and myotubes, including poly(A) RNA isolation, glyoxal/nitrite deamination, library preparation, and computational calling of m⁶A sites. **(B)** Phase-contrast images of proliferating and differentiated human myoblasts. Bar plots show RNA-seq expression levels (FPKM) of the myogenic marker *DESMIN* and the fibroblast marker *PDGFRA*, confirming myogenic identity and culture purity in both conditions. **(C)** Size-selected GLORI libraries following PAGE purification. Gel slices containing DNA fragments between 170–250 bp were excised and gel-purified for sequencing **(D)** Quantification of the total number of m⁶A sites detected in proliferating versus differentiated cells. **(E)** HOMER-derived consensus m⁶A sequence motif for proliferating and differentiated myoblasts. **(F)** Metagene distribution of GLORI-defined m⁶A sites across 5′UTR, CDS, and 3′UTR. **(G)** Distribution of per-site m⁶A stoichiometry in proliferating (top) and differentiated myoblasts (bottom). **(H)** IGV example showing single-nucleotide m⁶A sites in the 3′UTR of *H6PD*. A zoomed-in view highlights individual GLORI signals: orange marks A-to-G conversions (unmethylated adenosines), whereas green marks retained A bases (m⁶A-protected adenosines). This illustrates differential m⁶A presence and stoichiometry between proliferating and differentiated cells. **(I–J)** GO Biological Process enrichment analysis of genes containing m⁶A in proliferating (I) or differentiated (J) myoblasts. Both conditions share core functions such as Golgi-to-membrane protein transport and cytoskeleton regulation, while also displaying condition-specific signatures – including skeletal muscle differentiation, stress response and signalling pathways in proliferating cells, and mRNA modifications and calcium-entry pathways in differentiated cells – reflecting global shifts in the epitranscriptome during myogenic differentiation. Red highlighting indicates GO categories that are specific to each developmental stage.

As a positive control for method performance, we processed HEK293T mRNA in parallel, confirming robust and reproducible detection of m⁶A sites in agreement with the original GLORI report [31] (Supplementary Fig. 1A–E). Library quality assessment by native PAGE confirmed clean fragment distributions between 150–200 bp for all samples (Fig. 1C), indicating that RNA integrity was preserved throughout GLORI processing. Across human myogenic samples, we identified ∼70,000 high-confidence m⁶A sites in proliferating cells and ∼50,000 sites in differentiated myotubes mapping ∼8,000 expressed genes in proliferation and differentiation (Fig. 1D).

The marked reduction in modified sites upon differentiation echoes previously reported decreases in global m⁶A levels in mouse myoblast systems [16,21,30]. Notably, these studies reported substantially fewer m⁶A sites overall, likely reflecting both the more limited resolution of antibody-based approaches and potentially lower global m⁶A abundance in mouse muscle compared with human myogenic cells. By contrast, no global change was observed in a bovine differentiation model [32] highlighting species-specific regulation of the m⁶A epitranscriptome.

Motif analysis using HOMER revealed strong enrichment for the canonical DRACH consensus sequence, with the top motif matching GGACU, consistent across all samples (Fig. 1E). Metagene profiles showed characteristic accumulation of m⁶A around stop codons and within 3′ UTRs (Fig. 1F), in line with known m⁶A deposition patterns. Density analysis of per-site methylation levels further demonstrated an average of 46%, slightly higher than we see in HEK293T cells (42%) (Fig. 1G; Supplementary Fig. 1B). Inspection of individual sites in IGV confirmed these trends: for instance, a representative m⁶A site within the 3’UTR of the *HP6D* transcript displayed higher A→G conversion in differentiated cells than in proliferating cells, reflecting loss of methylation at this position (Fig 1H).

Finally, gene ontology analysis of m⁶A-modified transcripts in proliferating and differentiated myoblasts revealed both shared biological processes and state-specific functional enrichments, underscoring the dynamic and stage-dependent regulation of the m⁶A epitranscriptome during myogenesis (Fig 1I-J).

Together, these data demonstrate that m⁶A is remodelled during human skeletal muscle cell differentiation and is associated with pathways underlying myogenic biology. Importantly, our findings indicate that programs of m⁶A enrichment are associated with myoblast states, and that this regulation is not driven by large-scale changes in the positional distribution of m⁶A along transcripts.

### Distinct classes of regulated m⁶A sites are associated with proliferative and differentiated myogenic states

To explore how m⁶A methylation is remodeled during human myogenic differentiation, we first compared m⁶A-modified transcripts across proliferating and differentiated myoblasts. This analysis identified a large, shared core of methylated transcripts, together with cell-state–specific subsets, comprising approximately ∼1,300 proliferation-specific and ∼600 differentiation-specific m⁶A-modified genes (Fig. 2A). Genes selectively methylated in proliferating cells were enriched for biological functions related to DNA regulation and skeletal muscle embryonic programs, whereas differentiation-specific m⁶A-modified transcripts were preferentially associated with muscle maturation, contraction, and cell morphogenesis (Fig. 2B).

**Figure 2:**
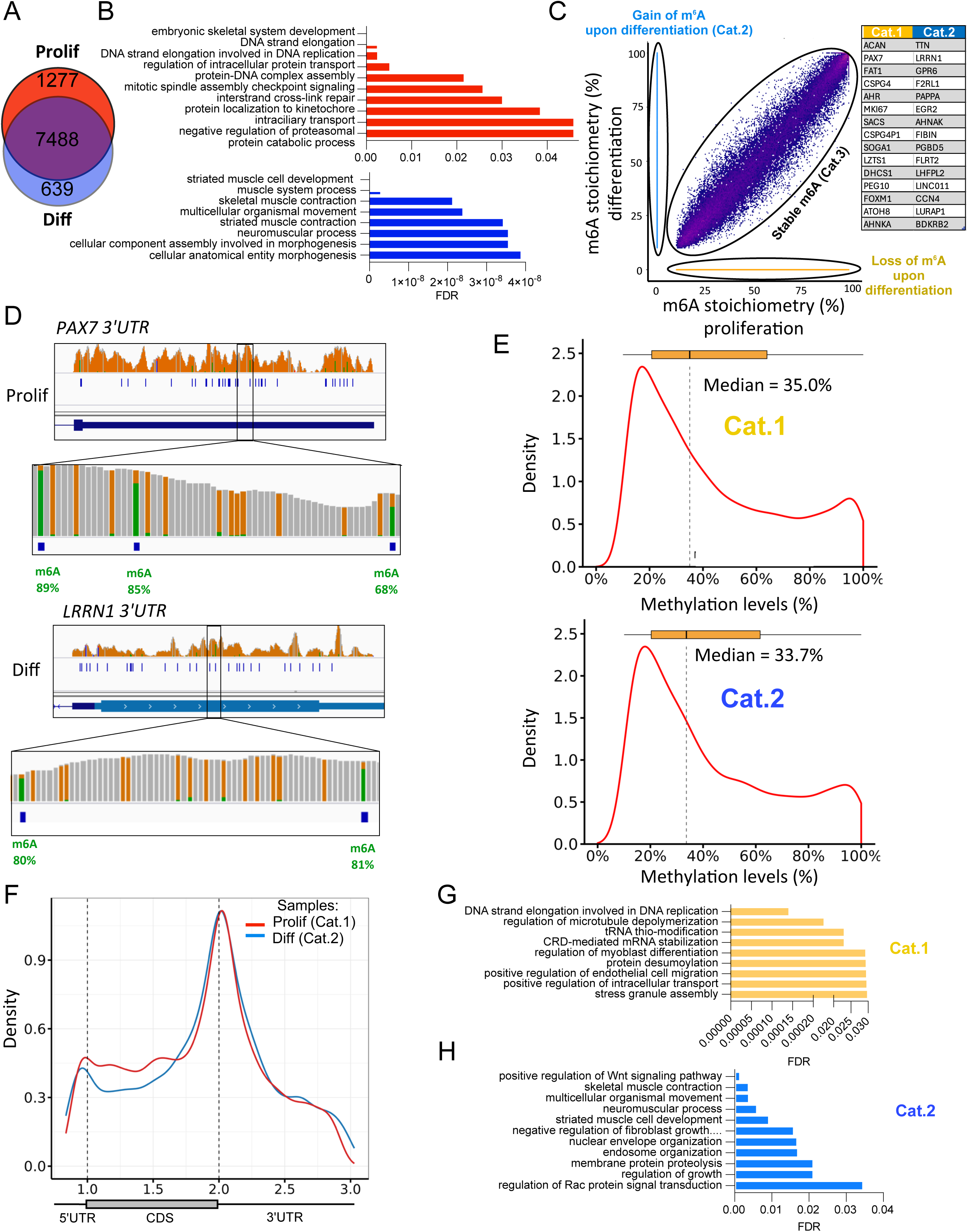
Differential m⁶A site usage during myogenic differentiation. **(A)** Overlap of m⁶A-containing genes between proliferating and differentiated myoblasts. **(B)** GO Biological Process enrichment analysis for genes uniquely methylated in proliferating (top) or differentiated (bottom) cells. **(C)** Scatter plot comparing per-site m⁶A levels in proliferating and differentiated myoblasts. Three major classes of regulated sites are highlighted: Category 1 (yellow), showing reduced m⁶A levels upon differentiation; Category 2 (blue), showing increased m⁶A levels upon differentiation; and Category 3 (diagonal), comprising sites with comparable methylation levels in proliferating and differentiated cells. Each dot represents an individual m⁶A site (apparent yellow and blue bands represent dense clusters of sites). Shared sites are visualized as a two-dimensional binned density distribution (color intensity reflects the number of sites per stoichiometry bin), whereas sites detected exclusively in one condition are projected onto the corresponding axis. The accompanying table lists representative genes exhibiting the strongest loss (left column) or gain (right colum) of m⁶A sites between conditions. **(D)** IGV examples of regulated m⁶A sites, showing sites localisation in *PAX7* (proliferation-specific methylation) and in *LRRN1* (differentiation-specific methylation). **(E)** Distributions of m⁶A stoichiometry in Class 1 and Class 2 sites. **(F)** Metagene profiles of m⁶A site distribution across the transcriptome (5′UTR, CDS, 3′UTR) for Cat. 1 (loss upon differentiation) and Cat. 2 (gain upon differentiation) m⁶A-regulated sites, highlighting differential m6A coverage. **(G–H)** GO Biological Process enrichment analysis for Cat. 1 (loss upon differentiation) and Cat. 2 (gain upon differentiation) m⁶A-regulated genes, revealing functions associated with cell cycle/DNA, and early-stage differentiation regulation (Cat. 1) and muscle maturation (Cat. 2).

To further resolve how m⁶A regulation operates at the nucleotide level, we next quantified per-site methylation stoichiometry between proliferating and differentiated myoblasts. This analysis was enabled using GLORI, a method introduced in 2023 that allows high-sensitivity, single-nucleotide detection and quantitative measurement of m⁶A stoichiometry – an approach that have not previously been applied to skeletal muscle cells.

This analysis revealed extensive and patterned epitranscriptomic remodeling during myogenic differentiation, with m⁶A sites segregating into three different regulatory classes (Fig. 2C). Class 1 sites displayed methylation in proliferating myoblasts but lost methylation upon differentiation, whereas Class 2 sites exhibited the opposite behavior, becoming selectively methylated in differentiated myotubes. These patterns indicate that m⁶A regulation during myogenesis does not result from uniform global shifts in methylation levels but instead arises from coordinated, site-specific modulation across distinct sets of transcripts. In contrast, class 3 sites lying near the diagonal exhibit similar methylation levels in proliferating and differentiated cells, reflecting only modest, proportionally scaled changes.

These likely represent intermediate or weakly responsive sites rather than strongly regulated classes. Together, this analysis suggests that the most functionally relevant m⁶A dynamics during myogenesis are concentrated within two subsets of class 1 and class 2 sites showing strong directional changes.

Inspection of representative loci using IGV confirmed these trends. For example, *PAX7* showed robust methylation sites in proliferating myoblasts but not in differentiated myotubes, whereas *LRRN1* displayed increased methylation upon differentiation (Fig. 2D)

Density analysis of site-specific methylation levels further revealed that differentially regulated m⁶A sites exhibited lower average methylation levels compared with the global m⁶A site population (Fig. 2E), consistent with these sites representing dynamic, actively remodeled loci rather than constitutively methylated regions. Metagene profiling showed that state-specific m⁶A sites retained the canonical enrichment around stop codons and within 3′ untranslated regions (3′UTRs) (Fig. 2F), indicating that global positional preferences are preserved. However, differences were observed within coding sequences (CDS) between proliferating and differentiated cells (Fig. 2F), suggesting condition-specific modulation of m⁶A levels within functional transcript regions.

Finally, gene ontology enrichment analysis of genes harboring state-specific m⁶A sites revealed engagement of more complex and specialized biological processes than those identified by transcript-level analyses alone, supporting a role for site-specific m⁶A regulation in shaping stage-dependent post-transcriptional programs during human myogenesis.

### The human myogenic m⁶A landscape is organized by site number, stoichiometry and expression

To further characterize transcript-level features of m⁶A regulation during human myogenesis, we examined the relationship between gene expression, m⁶A site number, and m⁶A stoichiometry in proliferating and differentiated myoblasts. This analysis indicates the presence of multiple regulatory programs, defined by distinct combinations of these parameters. One group of transcripts exhibited a high number of m⁶A sites together with moderate expression levels, reflecting extensive m⁶A marking across multiple positions. A second group comprised transcripts with a limited number of m⁶A sites but high m⁶A stoichiometry, consistent with strong modification at individual sites. A third category consisted of highly expressed transcripts carrying relatively few m⁶A sites. Notably, transcripts encoding muscle-related and regulatory factors were distributed across these different categories, and their relative positioning within this framework changed between proliferating and differentiated states. This redistribution highlights dynamic remodeling of the m⁶A landscape across myogenic differentiation and underscores the coexistence of multiple layers of m⁶A-associated transcript regulation within the myogenic program. Gene Ontology analysis of each category identified state-specific functional associations, with transcripts in proliferating and differentiated myoblasts enriched for different regulatory pathways (Fig. 3A–B).

**Figure 3:**
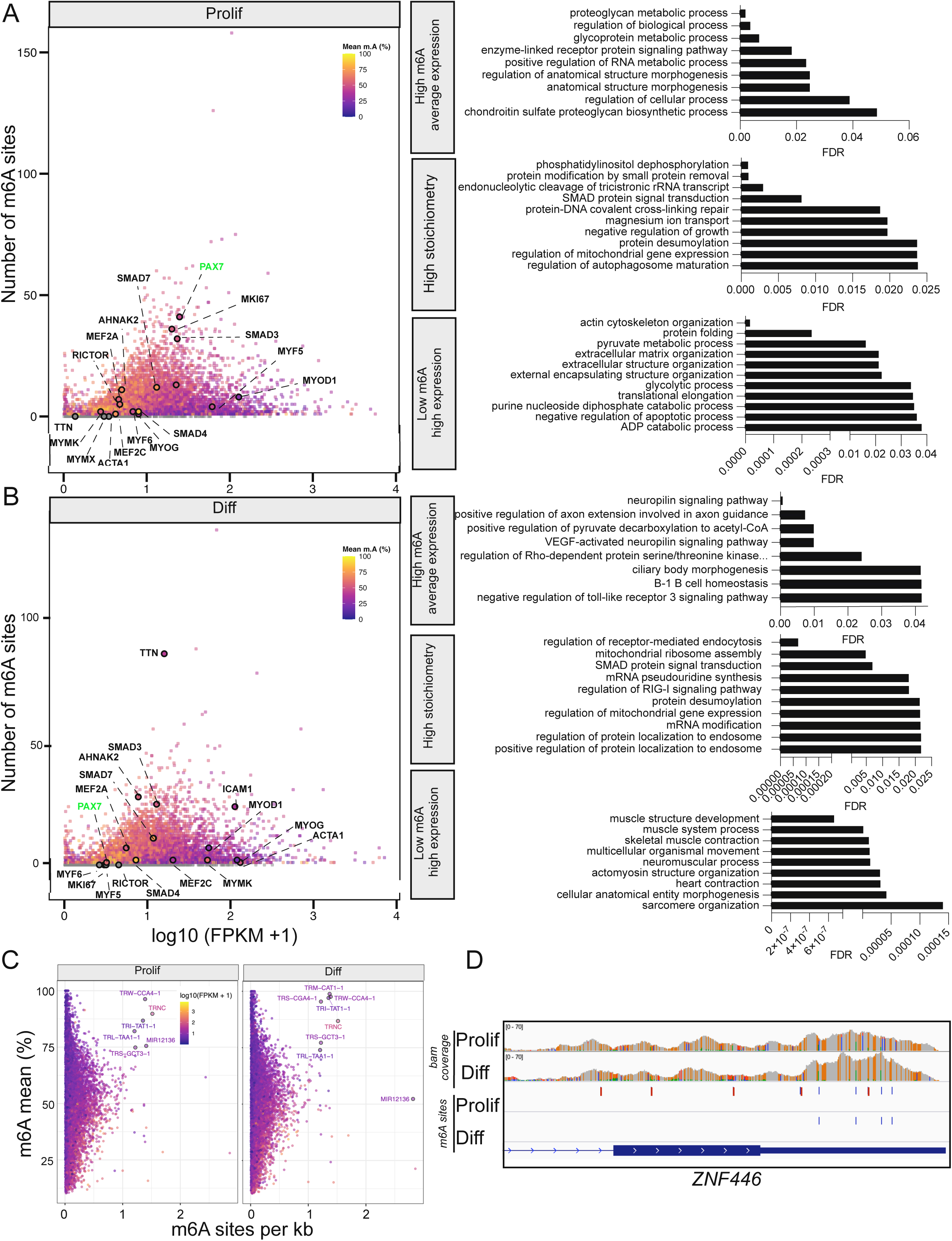
Expression, site number, and stoichiometry define transcript-level m⁶A organisation during human myogenesis. **(A–B)** Relationship between gene expression (log₁₀ FPKM+1) and the number of m⁶A sites per transcript in proliferating (A) and differentiated (B) myoblasts. Each point represents a transcript, coloured by mean m⁶A stoichiometry (%). Selected myogenic and regulatory genes are highlighted and annotated. Genes were stratified into three categories based on expression and m⁶A features: (i) high m⁶A site number with moderate expression (≥20 m⁶A sites; 0.5 ≤ log₁₀(FPKM + 1) ≤ 2.5), (ii) high m⁶A stoichiometry (0.5 ≤ log₁₀(FPKM + 1) ≤ 1), and (iii) low m⁶A / high expression (≤5 m⁶A sites; log₁₀(FPKM + 1) ≥2). For each category, Gene Ontology Biological Process enrichment analysis is shown on the right, ranked by false discovery rate (FDR). **(C)** Scatter plot showing per-gene m⁶A density (m⁶A sites per kb) versus mean methylation level (%) in proliferating (left) and differentiated (right) cells. Several tRNA genes appear as outliers with unusually high mean methylation despite modest m⁶A site density. **(D)** A representative IGV view of m⁶A sites across *ZNF446* in proliferating and differentiated cells. BAM read coverage is shown at the top, and GLORI-identified single-nucleotide m⁶A positions are shown below. Differentially methylated sites between conditions are highlighted in red.

Notably, transcripts with a high number of m⁶A sites did not necessarily display the highest mean methylation levels. Instead, genes harboring fewer m⁶A sites often exhibited larger changes in mean m⁶A stoichiometry between conditions, consistent with distinct regulatory strategies acting either through coordinated modulation of multiple sites or through strong regulation at a limited number of key positions (Fig. 3C). While such modes of m⁶A regulation have been proposed previously [33], our data provide direct single-nucleotide evidence for this phenomenon in human skeletal muscle cells. Inspection of representative transcripts, such as ZNF446, confirmed dynamic, site-specific m⁶A remodeling occurring without major changes in overall transcript abundance (Fig. 3D).

### Cross-species comparison and m⁶A reader association during human myogenic differentiation

Comparison of our human m⁶A dataset with a published mouse dataset identified both shared and human-specific m⁶A-modified genes and pathways, indicating a combination of common and species-specific epitranscriptomic regulation during myogenesis. In human proliferating myoblasts, species-specific m⁶A enrichment was observed in signalling and energy-regulation pathways, whereas in differentiated cells, human-specific signatures were predominantly associated with intracellular localization, transport and negative regulation of cell motility (Fig. 4A–B).

**Figure 4:**
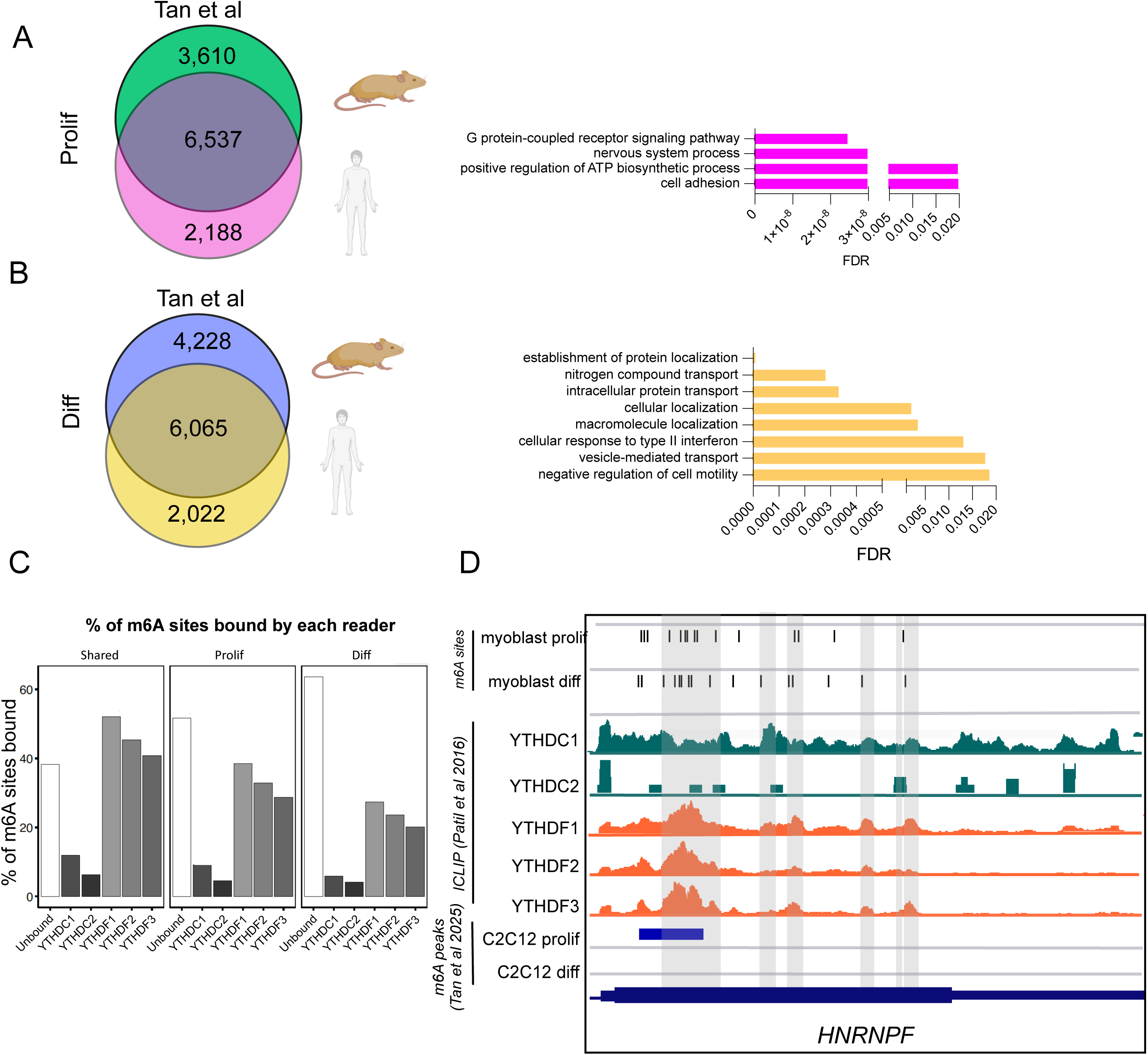
Cross-species conservation and reader recognition of m⁶A sites. **(A–B)** Cross-species comparison of m⁶A-modified genes based on the overlap between human GLORI datasets (Prolif or Diff) and mouse MeRIP-seq peaks (Tan *et al.*, 2025). Venn diagrams (left) show the shared and human-specific methylated genes, while GO Biological Process enrichment analysis (right) highlights functional differences: human-specific m⁶A genes in proliferating cells are enriched for signalling and adhesion pathways, whereas human-specific genes in differentiated cells are enriched for transport, localisation, and vesicle-mediated trafficking. **(C)** Public iCLIP datasets for YTHDF1/2/3 and YTHDC1/2 generated in HEK293T cells (Patil *et al.*, 2016 [29]) were intersected with GLORI single-nucleotide m⁶A sites to determine reader-associated sites. For each category (proliferation-only, differentiation-only, shared), bars indicate the percentage of m⁶A sites overlapping detectable reader signal — allowing multi-reader binding — as well as the fraction not overlapping any reader signal. **(D)** IGV tracks illustrating the relationship between GLORI-identified m⁶A sites and m⁶A reader binding profiles at the *HNRNPF* locus (displayed region corresponds to the last exon and portion of the 3′ UTR). Our human GLORI data (single-nucleotide m⁶A calls from proliferating and differentiated myoblasts) are shown together with iCLIP read distributions, as well as MeRIP-seq m⁶A peaks from mouse C2C12 myogenic cells (Tan et al., 2025). While GLORI identifies discrete, site-resolved m⁶A positions MeRIP-seq display broader enrichment patterns along transcripts.

Having established the presence of both conserved and human-specific m⁶A programs, we next examined how these sites are interpreted by m⁶A-binding proteins. Integration with published iCLIP datasets for m⁶A RNA-binding proteins from human cells (HEK293T) revealed differences in the proportion of diff-only, prolif-only, and shared sites bound by individual readers (Fig. 4C), pointing to distinct modes of reader engagement across myogenic states.

Visual inspection of representative loci further confirmed concordance between dynamically regulated, site-resolved m⁶A positions identified by GLORI and local enrichment of reader-binding signal (Fig. 4D), supporting the functional relevance of single-nucleotide m⁶A remodeling. By contrast, antibody-based MeRIP-seq approaches often lack the sensitivity and resolution required to detect discrete or condition-specific m⁶A sites, leading to undetected or poorly resolved events—such as those observed at the *HNRNPF* locus.

## Discussion

Skeletal muscle is the most abundant tissue in the human body and plays vital roles far beyond locomotion. It is essential for respiration, energy expenditure, glucose, amino acid, and lipid metabolisms, as well as overall quality of life [34,35]. Following injury, skeletal muscle activates a highly coordinated regeneration program in which muscle stem cells and their niche collaborate to restore tissue structure and function. This regenerative response is remarkably efficient, but its precision also makes it vulnerable: even subtle impairments can lead to profound deficits and contribute to a wide spectrum of debilitating disorders [1]. Given these complex functions, the mechanisms governing muscle physiology must be regulated with exceptional accuracy.

Skeletal muscle differentiation is a highly dynamic, time-dependent process characterized by coordinated transitions from stem cell quiescence to activation, proliferation, and terminal differentiation [36]. These tightly regulated steps underpin skeletal muscle regeneration. To date, research has primarily focused on transcriptional and signaling pathways governing myogenic progression [37]. However, comparatively little is known about how post-transcriptional mechanisms—particularly RNA modifications—contribute to the control of myogenesis [4,37,38]. This gap is striking given the growing recognition that the epitranscriptome represents a critical regulatory layer of gene expression. More than 170 RNA modifications have been identified, with RNA methylation emerging as the most abundant and functionally influential class. Among these, N⁶-methyladenosine (m⁶A) has been implicated in diverse processes including RNA stability, translation, and cell fate decisions across multiple developmental contexts. Despite this, an important unresolved question is how RNA modifications are selectively deployed to ensure precise temporal control of myogenic gene expression programs.

In the experiments described here, we provide the first evidence that m⁶A methylation in human myogenic cells undergoes extensive remodeling at the level of individual nucleotides during differentiation. Importantly, these changes are not reflected by large shifts in global methylation levels; instead, they mostly manifest as highly selective, site-specific gains and losses of m⁶A on transcripts with key roles in muscle biology. To achieve this level of precision, we used GLORI, a recently developed chemical-based method that enables absolute quantification of m⁶A at single-nucleotide resolution without relying on antibodies. This technique overcomes key limitations of earlier approaches - such as low resolution, background noise, and bias - making it uniquely suited to detect subtle but functionally meaningful remodeling events during myogenesis.

To benchmark the robustness and reproducibility of our approach, we profiled HEK293T cells as an external reference. We observed a very high concordance with the original GLORI study, demonstrating not only our ability to perform this technically challenging methodology, but also the strong reproducibility of GLORI across laboratories.

Consistent with previous reports in other cell types and tissues, we detected tens of thousands of m⁶A sites genome-wide in human cells. Moreover, the average m⁶A motif composition and transcript body distribution closely matched what has been described so far, with enrichment around stop codons and 3’UTR, indicating that the overall architecture of the m⁶A epitranscriptome is conserved in human muscle cells [7,39].

We found that m⁶A methylation is dynamically regulated during myogenic differentiation. Pathways associated with regulated m⁶A sites were biologically relevant for each cellular state, supporting a functional role for m⁶A remodeling in the transition between proliferating myoblasts and differentiated myotubes.

Notably, our data reveal dynamic m⁶A regulation at specific sites on canonical myogenic regulators, including PAX7, MYOD1, and MYOG. This contrasts with previous reports – for example, those by Gheller et al. [16] which did not detect m⁶A changes on these factors during differentiation in mouse myogenic cells. When comparing our dataset with mouse MeRIP-seq profiles, we identified human-specific epitranscriptomic signatures, including pathways and individual methylation sites present in human myogenic cells but absent in mice. This discrepancy underscores the complexity of the m⁶A landscape and highlights the importance of species-specific analyses. Whereas earlier approaches primarily captured transcript- or region-level methylation, our single-nucleotide mapping strategy reveals fine-scale remodeling events that would likely remain undetected using lower-resolution methodologies.

Importantly, the observed differences cannot be attributed solely to species-related effects, as they may also be influenced by the use of immortalized C2C12 cells in mouse studies, which may differ from primary myogenic cells in their epitranscriptomic regulation. A direct comparison using primary mouse myoblasts would be required to disentangle species-specific from cell-intrinsic effects; however, to the best of our knowledge, such single-nucleotide m⁶A profiling has not yet been performed in this context. Therefore, we cannot exclude that such cell-intrinsic effects act in parallel with, or account for, the apparent species-specific differences observed.

A particularly striking observation emerged when examining m⁶A sites unique to one cellular state. Stage-specific m⁶A sites exhibited distinct positional distributions along the transcript body and displayed globally reduced average methylation levels, consistent with selective – not uniform – m⁶A regulation. Gene Ontology analyses of these stage-restricted sites highlighted pathways tightly aligned with the biological processes governing each differentiation stage, pointing to fine-tuned epitranscriptomic control of myogenic progression.

To further integrate these findings at the transcript level, we classified m⁶A-modified genes into three broad categories based on the relationship between transcript abundance, number of m⁶A sites, and mean methylation stoichiometry. This analysis revealed that myogenic regulators are not confined to a single epitranscriptomic regime but are instead distributed across all three categories. Some transcripts harbor multiple m⁶A sites with moderate stoichiometry, suggesting coordinated regulation through cumulative modification, whereas others carry only one or a few sites that exhibit high methylation levels, consistent with strong regulation at strategically positioned nucleotides. A third group comprises highly expressed transcripts with relatively sparse m⁶A marking, indicating a more permissive mode of regulation in which m⁶A likely modulate RNA fate rather than acting as a dominant regulatory switch. The presence of canonical myogenic factors across these distinct categories argues against a uniform mode of m⁶A action and instead supports a model in which m⁶A operates through multiple, parallel regulatory strategies. In this framework, myogenic transcripts act as hubs for layered post-transcriptional control, where site number, stoichiometry, and reader engagement collectively shape gene-specific regulatory outcomes during myogenic differentiation.

These observations emphasize that the absence of detectable changes at the global or gene-body level does not preclude functionally meaningful regulation at specific, strategically positioned sites.

Finally, because the function of m⁶A depends on the proteins that recognize it, we examined the behavior of m⁶A “reader” proteins. Readers such as YTHDF1–3, YTHDC1-2 family members bind methylated sites and determine their fate – promoting splicing, enhancing stability, or regulating decay. Our single-nucleotide maps revealed that differentiation-associated m⁶A remodeling occurs at sites bound by specific readers, and that distinct classes of readers preferentially associate with proliferative versus differentiation-specific methylation patterns. This suggests that the biological outcome of m⁶A remodeling is mediated not only by which sites are methylated, but also by which reader proteins are available to interpret these changes in each cellular state.

## Conclusions

In this study, we investigate the epitranscriptome for the first time during human myogenesis. We applied the recently developed GLORI approach—based on glyoxal and nitrite-mediated deamination of unmethylated adenosines—which enables single-nucleotide–resolution and stoichiometry-resolved mapping of m⁶A in primary human myoblasts and their differentiated counterparts. Our analyses demonstrate the high quality and robustness of the datasets and reveal a global decrease in m⁶A levels during myogenic differentiation. This reduction is observed both at the gene level and at the level of individual m⁶A sites and their methylation stoichiometry, indicating a complex, multi-layered regulation of the m⁶A epitranscriptome during myogenesis. m⁶A mapping highlights dynamic regulation of transcripts encoding key myogenic regulators and pathways. Importantly, normalization to gene expression levels shows that these changes are not solely driven by transcriptional variation, supporting an independent layer of post-transcriptional control. Comparison with published mouse m⁶A datasets during differentiation uncovers a human-specific m⁶A signature. Finally, a subset of these sites overlaps with binding sites of distinct m⁶A RNA-binding proteins, revealing an additional layer of regulation mediated by reader-specific engagement and supporting state-dependent interpretation of m⁶A marks. Our findings strongly support this model in skeletal muscle, demonstrating that myogenic differentiation is accompanied by precise reprogramming of the m⁶A epitranscriptome rather than uniform methylation shifts. Taken together, this work positions m⁶A as a critical layer of post-transcriptional regulation in human muscle differentiation and provides a conceptual framework for understanding how RNA modifications contribute to muscle plasticity in both physiological and pathological contexts.

## List of abbreviations

m⁶A: N⁶-methyladenosine
GLORI: glyoxal- and nitrite-mediated deamination of unmethylated adenosines
RBP: RNA-binding protein
CLIP-seq: crosslinking and immunoprecipitation sequencing
MeRIP-seq: methylated RNA immunoprecipitation sequencing
RNA-seq: RNA sequencing
UMI: unique molecular identifier
IGV: Integrative Genomics Viewer
GO: Gene Ontology
FDR: false discovery rate
uTPM: unique transcripts per million
FPKM: fragments per kilobase of transcript per million mapped reads
UTR: untranslated region
CDS: coding sequence
PAGE: polyacrylamide gel electrophoresis
PBS: phosphate-buffered saline
DMEM: Dulbecco’s modified Eagle’s medium
FBS: fetal bovine serum
EDTA: ethylenediaminetetraacetic acid
DMSO: dimethyl sulfoxide
MES: 2-(N-morpholino)ethanesulfonic acid
TEAA: triethylammonium acetate
TE: Tris–EDTA
PCR: polymerase chain reaction
HPC: high-performance computing
IFB: Institut Français de Bioinformatique
ANR: Agence Nationale de la Recherche
MSCA: Marie Skłodowska-Curie Actions
GEO: Gene Expression Omnibus

## Declarations

### Ethic approval and consent to participate

Not applicable

### Consent for publication

Not applicable

### Availability of data and materials

The datasets generated and/or analysed during the current study are available in the GEO repository, https://www.ncbi.nlm.nih.gov/geo/query/acc.cgi?acc=GSE78030, https://www.ncbi.nlm.nih.gov/geo/query/acc.cgi?acc=GSE293212

### Authors’ contributions

P.K. conceived, designed the study, performed GLORI-seq experiments, analysed the data, interpreted the results, and wrote the manuscript.

K.Z. performed cell culture experiments under the supervision of L.A. F.R provided support and input to the project. D.F. contributed to project supervision

## Acknowledgments

The authors thank Fiorella Grandi for her insightful feedback on the manuscript and valuable discussions. They are grateful to Nami Altin for providing the HEK293T cells. The authors also acknowledge the Myoline platform at the Institut de Myologie for providing the cell line, and the sequencing facility at the Institut Pasteur for performing the sequencing.

---

**Figure S1:**
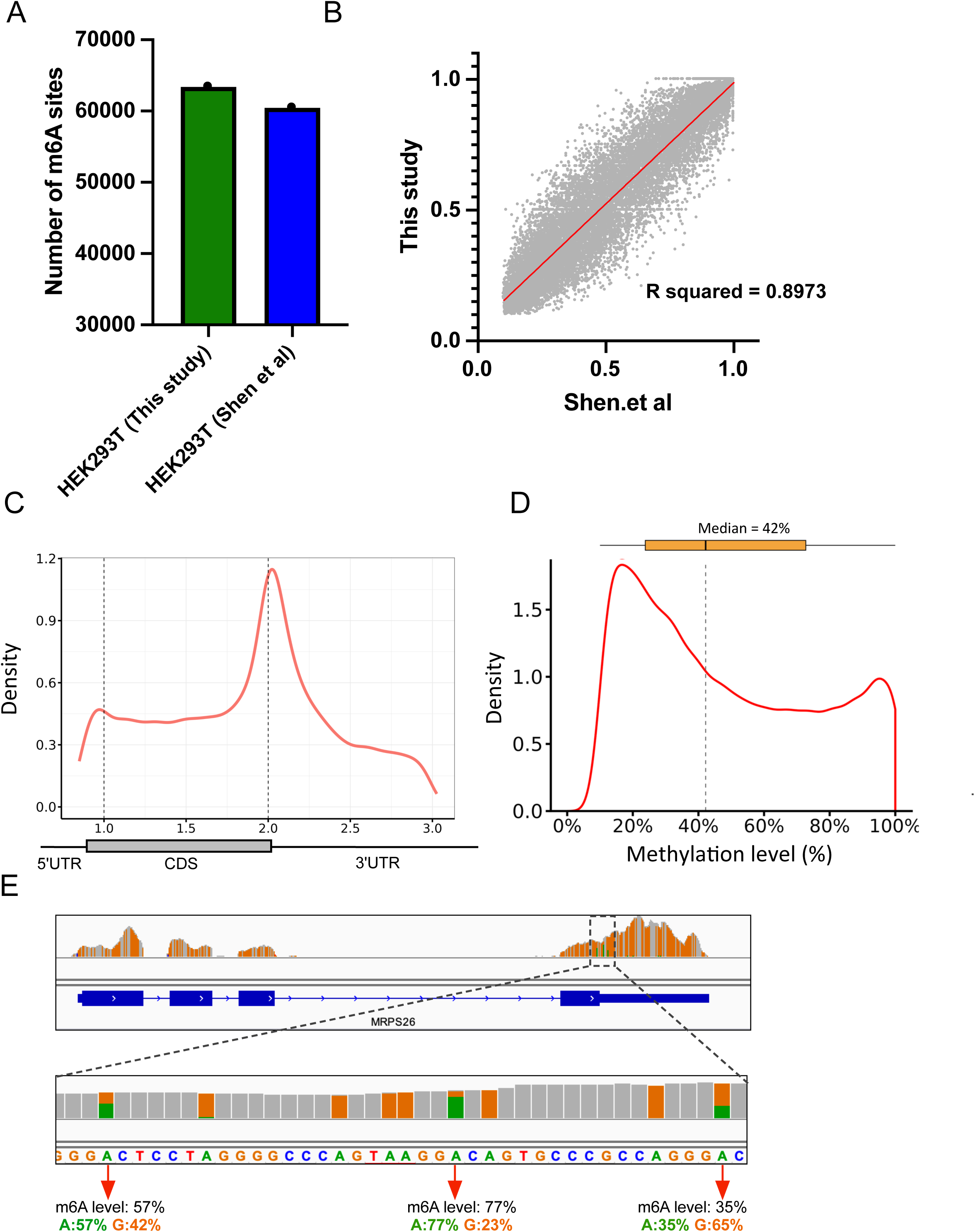
Benchmarking GLORI in HEK293T cells against published m⁶A GLORI reference dataset. (A) Total number of m⁶A sites identified by GLORI in HEK293T cells from this study and in those reported by *Shen et al.* ([31]). (B) Correlation of m⁶A stoichiometry values at shared sites (42073) between HEK293T cells analysed in this study and those reported by Shen *et al.*. Each dot represents an individual m⁶A site detected in both datasets. The coefficient of determination (R²) is indicated on the plot. (C) Metagene analysis showing the distribution of m⁶A sites along transcript regions (5′UTR, CDS, and 3′UTR). (D) Density distribution of m⁶A methylation levels (%) across all identified sites. The dashed line indicates the median methylation level. Value is reported on the plot. (E) Representative IGV view illustrating GLORI-identified single-nucleotide m⁶A sites across the *MRPS26* locus. A zoomed-in view highlights individual GLORI signals, where orange marks A-to-G conversions corresponding to unmethylated adenosines, and green marks retained adenosines protected from deamination, corresponding to m⁶A-modified sites, thereby illustrating site-specific methylation stoichiometry.

## References

1. Mukund K, Subramaniam S. Skeletal muscle: A review of molecular structure and function, in health and disease. Wiley Interdiscip Rev Syst Biol Med [Internet]. 2020;12:e1462. Available from: 10.1002/wsbm.1462

2. Martin PT. Role of transcription factors in skeletal muscle and the potential for pharmacological manipulation. Curr Opin Pharmacol [Internet]. 2003;3:300–8. Available from: 10.1016/s1471-4892(03)00050-x

3. Shi D-L, Grifone R. RNA-binding proteins in the post-transcriptional control of skeletal muscle development, regeneration and disease. Front Cell Dev Biol [Internet]. 2021;9:738978. Available from: 10.3389/fcell.2021.738978

4. Weskamp K, Olwin BB, Parker R. Post-transcriptional regulation in skeletal muscle development, repair, and disease. Trends Mol Med [Internet]. 2021;27:469–81. Available from: 10.1016/j.molmed.2020.12.002

5. Cappannini A, Ray A, Purta E, Mukherjee S, Boccaletto P, Moafinejad SN, et al. MODOMICS: a database of RNA modifications and related information. 2023 update. Nucleic Acids Res [Internet]. 2024;52:D239–44. Available from: 10.1093/nar/gkad1083

6. Liu C, Sun H, Yi Y, Shen W, Li K, Xiao Y, et al. Absolute quantification of single-base m6A methylation in the mammalian transcriptome using GLORI. Nat Biotechnol [Internet]. 2023;41:355–66. Available from: 10.1038/s41587-022-01487-9

7. Zaccara S, Ries RJ, Jaffrey SR. Reading, writing and erasing mRNA methylation. Nat Rev Mol Cell Biol [Internet]. 2019;20:608–24. Available from: 10.1038/s41580-019-0168-5

8. Geula S, Moshitch-Moshkovitz S, Dominissini D, Mansour AA, Kol N, Salmon-Divon M, et al. m 6 A mRNA methylation facilitates resolution of naïve pluripotency toward differentiation. Science [Internet]. 2015;347:1002–6. Available from: 10.1126/science.1261417

9. Batista PJ, Molinie B, Wang J, Qu K, Zhang J, Li L, et al. M6A RNA modification controls cell fate transition in mammalian embryonic stem cells. Cell Stem Cell [Internet]. 2014;15:707–19. Available from: 10.1016/j.stem.2014.09.019

10. Wang Y, Li Y, Toth JI, Petroski MD, Zhang Z, Zhao JC. N6-methyladenosine modification destabilizes developmental regulators in embryonic stem cells. Nat Cell Biol [Internet]. 2014;16:191–8. Available from: 10.1038/ncb2902

11. Wen J, Lv R, Ma H, Shen H, He C, Wang J, et al. Zc3h13 regulates nuclear RNA m6A methylation and mouse embryonic stem cell self-renewal. Mol Cell [Internet]. 2018;69:1028–38.e6. Available from: 10.1016/j.molcel.2018.02.015

12. Zhang C, Chen Y, Sun B, Wang L, Yang Y, Ma D, et al. m6A modulates haematopoietic stem and progenitor cell specification. Nature [Internet]. 2017;549:273–6. Available from: 10.1038/nature23883

13. Lv J, Zhang Y, Gao S, Zhang C, Chen Y, Li W, et al. Endothelial-specific m6A modulates mouse hematopoietic stem and progenitor cell development via Notch signaling. Cell Res [Internet]. 2018;28:249–52. Available from: 10.1038/cr.2017.143

14. Li L, Zang L, Zhang F, Chen J, Shen H, Shu L, et al. Fat mass and obesity-associated (FTO) protein regulates adult neurogenesis. Hum Mol Genet [Internet]. 2017;26:2398–411. Available from: 10.1093/hmg/ddx128

15. Klein P, Petrić Howe M, Harley J, Crook H, Esteban Serna S, Roumeliotis TI, et al. m6a methylation orchestrates IMP1 regulation of microtubules during human neuronal differentiation. Nat Commun [Internet]. 2024;15:4819. Available from: 10.1038/s41467-024-49139-7

16. Gheller BJ, Blum JE, Fong EHH, Malysheva OV, Cosgrove BD, Thalacker-Mercer AE. A defined N6-methyladenosine (m6A) profile conferred by METTL3 regulates muscle stem cell/myoblast state transitions. Cell Death Discov [Internet]. 2020;6:95. Available from: 10.1038/s41420-020-00328-5

17. Liang Y, Han H, Xiong Q, Yang C, Wang L, Ma J, et al. METTL3-mediated m6A methylation regulates muscle stem cells and muscle regeneration by Notch signaling pathway. Stem Cells Int [Internet]. 2021;2021:9955691. Available from: 10.1155/2021/9955691

18. Liu J, Zuo H, Wang Z, Wang W, Qian X, Xie Y, et al. The m6A reader YTHDC1 regulates muscle stem cell proliferation via PI4K-Akt-mTOR signalling. Cell Prolif [Internet]. 2023;56:e13410. Available from: 10.1111/cpr.13410

19. Qiao Y, Sun Q, Chen X, He L, Wang D, Su R, et al. Nuclear m6A reader YTHDC1 promotes muscle stem cell activation/proliferation by regulating mRNA splicing and nuclear export. Elife [Internet]. 2023;12. Available from: 10.7554/eLife.82703

20. Wang X, Huang N, Yang M, Wei D, Tai H, Han X, et al. FTO is required for myogenesis by positively regulating mTOR-PGC-1α pathway-mediated mitochondria biogenesis. Cell Death Dis [Internet]. 2017;8:e2702–e2702. Available from: 10.1038/cddis.2017.122

21. Xie S-J, Lei H, Yang B, Diao L-T, Liao J-Y, He J-H, et al. Dynamic m6A mRNA methylation reveals the role of METTL3/14-m6A-MNK2-ERK signaling axis in skeletal muscle differentiation and regeneration. Front Cell Dev Biol [Internet]. 2021;9:744171. Available from: 10.3389/fcell.2021.744171

22. Decary S, Mouly V, Hamida CB, Sautet A, Barbet JP, Butler-Browne GS. Replicative potential and telomere length in human skeletal muscle: implications for satellite cell-mediated gene therapy. Hum Gene Ther [Internet]. 1997;8:1429–38. Available from: 10.1089/hum.1997.8.12-1429

23. González MN, de Mello W, Butler-Browne GS, Silva-Barbosa SD, Mouly V, Savino W, et al. HGF potentiates extracellular matrix-driven migration of human myoblasts: involvement of matrix metalloproteinases and MAPK/ERK pathway. Skelet Muscle [Internet]. 2017;7:20. Available from: 10.1186/s13395-017-0138-6

24. Olarerin-George AO, Jaffrey SR. MetaPlotR: a Perl/R pipeline for plotting metagenes of nucleotide modifications and other transcriptomic sites. Bioinformatics [Internet]. 2017;33:1563–4. Available from: 10.1093/bioinformatics/btx002

25. Bailey TL, Boden M, Buske FA, Frith M, Grant CE, Clementi L, et al. MEME SUITE: tools for motif discovery and searching. Nucleic Acids Res [Internet]. 2009;37:W202–8. Available from: 10.1093/nar/gkp335

26. Mi H, Ebert D, Muruganujan A, Mills C, Albou L-P, Mushayamaha T, et al. PANTHER version 16: a revised family classification, tree-based classification tool, enhancer regions and extensive API. Nucleic Acids Res [Internet]. 2021;49:D394–403. Available from: 10.1093/nar/gkaa1106

27. Feuermann M, Mi H, Gaudet P, Muruganujan A, Lewis SE, Ebert D, et al. A compendium of human gene functions derived from evolutionary modelling. Nature [Internet]. 2025;640:146–54. Available from: 10.1038/s41586-025-08592-0

28. Supek F, Bošnjak M, Škunca N, Šmuc T. REVIGO summarizes and visualizes long lists of gene ontology terms. PLoS One [Internet]. 2011;6:e21800. Available from: 10.1371/journal.pone.0021800

29. Patil DP, Chen C-K, Pickering BF, Chow A, Jackson C, Guttman M, et al. m6A RNA methylation promotes XIST-mediated transcriptional repression. Nature [Internet]. 2016;537:369–73. Available from: 10.1038/nature19342

30. Tan Y-Y, Ou Y-W, Zuo Q, Luo Y, Chen W-C, Chen W-X, et al. Stage-specific requirement for METTL3-dependent m6A epitranscriptomic regulation during myogenesis. Commun Biol [Internet]. 2025;8:1317. Available from: 10.1038/s42003-025-08759-5

31. Shen W, Sun H, Liu C, Yi Y, Hou Y, Xiao Y, et al. GLORI for absolute quantification of transcriptome-wide m6A at single-base resolution. Nat Protoc [Internet]. 2024;19:1252–87. Available from: 10.1038/s41596-023-00937-1

32. Yang X, Wang J, Ma X, Du J, Mei C, Zan L. Transcriptome-wide N 6-methyladenosine methylome profiling reveals m6A regulation of skeletal myoblast differentiation in cattle (Bos taurus). Front Cell Dev Biol [Internet]. 2021;9:785380. Available from: 10.3389/fcell.2021.785380

33. Murakami S, Jaffrey SR. Hidden codes in mRNA: Control of gene expression by m6A. Mol Cell [Internet]. 2022;82:2236–51. Available from: 10.1016/j.molcel.2022.05.029

34. Baskin KK, Winders BR, Olson EN. Muscle as a “mediator” of systemic metabolism. Cell Metab [Internet]. 2015;21:237–48. Available from: 10.1016/j.cmet.2014.12.021

35. Sartori R, Romanello V, Sandri M. Mechanisms of muscle atrophy and hypertrophy: implications in health and disease. Nat Commun [Internet]. 2021;12:330. Available from: 10.1038/s41467-020-20123-1

36. Yin H, Price F, Rudnicki MA. Satellite cells and the muscle stem cell niche. Physiol Rev [Internet]. 2013;93:23–67. Available from: 10.1152/physrev.00043.2011

37. Relaix F, Bencze M, Borok MJ, Der Vartanian A, Gattazzo F, Mademtzoglou D, et al. Perspectives on skeletal muscle stem cells. Nat Commun [Internet]. 2021;12. Available from: 10.1038/s41467-020-20760-6

38. Imbriano C, Moresi V, Belluti S, Renzini A, Cavioli G, Maretti E, et al. Epitranscriptomics as a New Layer of Regulation of Gene Expression in Skeletal Muscle: Known Functions and Future Perspectives. Int J Mol Sci [Internet]. 2023;24. Available from: 10.3390/ijms242015161

39. Meyer KD, Saletore Y, Zumbo P, Elemento O, Mason CE, Jaffrey SR. Comprehensive analysis of mRNA methylation reveals enrichment in 3’ UTRs and near stop codons. Cell [Internet]. 2012;149:1635–46. Available from: 10.1016/j.cell.2012.05.003

